# Interaction between SARS-CoV-2 spike glycoprotein and human skin models: a molecular dynamics study

**DOI:** 10.1101/2021.07.13.452154

**Authors:** Marc Domingo, Jordi Faraudo

## Abstract

The possibility of contamination of human skin by infectious virions plays an important role in indirect transmission of respiratory viruses but little is known about the fundamental physico-chemical aspects of the virus-skin interactions. In the case of coronaviruses, the interaction with surfaces (including the skin surface) is mediated by their large glycoprotein spikes that protrude from (and cover) the viral envelope. Here, we perform all atomic simulations between the SARS-CoV-2 spike glycoprotein and human skin models. We consider an “oily” skin covered by sebum and a “clean” skin exposing the stratum corneum. The simulations show that the spike tries to maximize the contacts with stratum corneum lipids, particularly ceramides, with substantial hydrogen bonding. In the case of “oily” skin, the spike is able to retain its structure, orientation and hydration over sebum with little interaction with sebum components. Comparison of these results with our previous simulations of the interaction of SARS-CoV-2 spike with hydrophilic and hydrophobic solid surfaces, suggests that the”soft” or “hard” nature of the surface plays an essential role in the interaction of the spike protein with materials.

## 1 Introduction

The novel coronavirus SARS-CoV-2 emerged in December 2019 as a human pathogen^1^ that caused the COVID-19 disease world pandemic^2^. It is the third documented spillover of an animal coronavirus to humans in only two decades that has resulted in a major epidemic^1^ (SARS-CoV, MERS and SARS-CoV-2). Prior to the emergence of SARS-CoV in 2002, only two human coronaviruses (HCoV-OC43 and HCoV-229E) were known, which cause mild respiratory tract infections (including the common cold). Other important coronaviruses pathogenic to humans include^3^ HCoV-NL63 (discovered in 2004) that causes mild respiratory infection and HCoV-HKU1 (discovered in 2005) that causes pneumonia. Therefore, it is clear that emerging infectious diseases caused by coronaviruses must be seen as a major threat to human health.

Coronaviruses belong to the category of enveloped viruses, which means that they have a membrane (envelope)^4,5^. In the case of a coronavirus virion, the major components of the envelope^6^ are the transmembrane matrix glycoprotein (M) and lipids that the virion obtained from the host cell ^5^. But the most prominent feature of coronaviruses is the presence of a large glycoprotein spike (S) protruding from the envelope which give a characteristic appearance to this virus family and give them they name (from “corona” which is the latin for “crown”). The S protein is responsible for the interaction of the virus with a host cell receptor.

The transmission of respiratory viruses in general (including coronaviruses and SARS-CoV-2 in particular) involves the expiratory emission of virus-containing aerosols and droplets^7,8^ which may infect other individuals via direct or indirect mechanisms. The direct mechanism involves inhalation of aerosols or the deposition of emitted droplets on mucosal surfaces (e.g. mouth, eyes). Indirect transmission occurs through physical contact with virus containing aerosols and droplets deposited onto materials (exposed surfaces of common objects such as furniture or electronic gadgets, textiles, protective equipment,…) and byself inoculation of virus into the mouth, nose or eye as illustrated in Figure 1. Another indirect source of transmission is reaerosolization after deposition onto these surfaces.

**Fig. 1.**
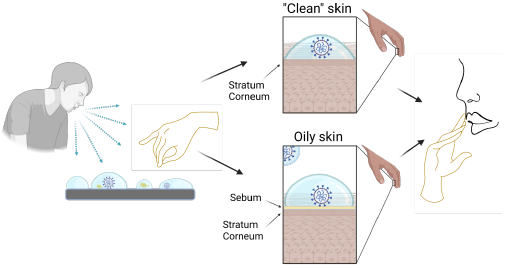
Scheme of the role of skin in the indirect transmission of a virus particle via contaminated surfaces. The sneezing or coughing of an infected individual can contaminate a surface or directly the skin of another individual. The virion may remain intact and infectious over the skin (either a sebaceous, oily skin or a clean skin with the stratum corneum exposed) so the individual can be then infected by touching their eyes mouth or nose. Scheme created with BioRender.com

Indirect transmission has been clearly demonstrated by previous work in different enveloped respiratory viruses ^7^. For example, empirical studies ^9^ following daily life activity of persons infected with common cold demonstrate environmental contamination with rhinovirus at surfaces of door handles, pens, light switches and subsequent transfer to fingers of healthy individuals. Other studies ^7^ found infectious influenza viruses in the air after shaking of a contaminated blanket, from resuspension of contaminated dust or even viruses aerosolized from paper tissues previously contaminated by an infected individual.

Coronavirus particles are relatively robust, and deposited virions remain infectious for several days when adsorbed onto a variety of materials even in relatively harsh conditions^3,10,11^. This stability of the adsorbed coronavirus particle suggests that these indirect mechanisms described above may have a significant relevance in contributing to coronavirus spread^3^ as emphasized by the WHO organisation^8^. In fact, the persistence of viable virus onto surfaces is the reason for the recommendation of health authorities worldwide on continually disinfecting and cleaning surfaces and washing hands.

In this respect, it is clear that the interaction of human skin with virus-containing droplets plays a role in indirect transmission of viruses, as emphasized for example in Ref ^12^. The importance of the indirect mode of transmission indicated schematically in Figure 1 is expected to depend critically on factors such as the adhesion strength of a virion over human skin and whether the virion particle retains or not its integrity and its hydration upon adhesion onto human skin.

Interestingly, experimental studies ^13^ show that the SARS-CoV-2 virus is able to survive infectious adsorbed over human skin for about 10 hours, much longer than typical survival time for influenza virus (∼ 1 − 2 hours). These studies also show that SARS-CoV-2 virus adsorbed onto skin are inactivated within 15 seconds upon treatment with ethanol based disinfectants, emphasizing the importance of correct hand hygiene protocols.

The reason for this long stability of the SARS-CoV-2 virus over human skin is not know. In fact, at the present time, there is a lack of fundamental understanding of fundamental aspects of the interactions between coronaviruses and surfaces at the physicochemical level. This is in fact the main motivation for this paper: to elucidate the molecular details of the interaction between SARS-CoV-2 virus and human skin. This study will also complement our previous work on the interaction between SARS-CoV-2 and different materials^14^.

In the case of coronaviruses, the interaction of the virus with the environment takes place through the large spike protein S protruding from the virus envelope^4^, since the envelope is densely covered by spikes (∼ 100 spike copies for a ∼ 100 nm virion ^15^ in the case of SARS-CoV-2). Given the fact that the molecular structure and atomistic coordinates of the S protein are known^16^, we will consider here atomistic molecular dynamics simulations of the interaction between S and a model of the external surface of the human skin.

Of course, modelling the surface of skin is a problem of great complexity, so appropriate simplifications are needed. We will distinguish between two different situations (see Figure 1). One case corresponds to clean skin in body regions without hair and without sebaceous glands, such as the palms of the hands. In this case, the exposed surface of the skin corresponds to the outermost layer of the epidermis, the so called stratum corneum (SC)^17^. The opposite case corresponds to skin covered by a protective oily or waxy layer known as sebum^18^, which is segregated by the sebaceous glands, which are present in large numbers in our face, for example. Sebum can be present not only in regions of the skin with sebaceous glands but also in other body parts (for example fingers) in which sebum has been transferred by touching other skin regions covered by sebum. In this case, sebum can be easily removed by using personal care products such as soap.

Therefore, we will consider the interaction of the spike protein S of the SARS-CoV-2 virus with both a model of the SC and a model of a sebum layer. These two extreme and somewhat simplified situations (clean SC without sebum and skin covered by sebum) are indicated as “clean” skin and oily skin respectively in Figure 1.

It is interesting to remark that the surfaces of the SC and the sebum have very different molecular organization and properties (for example regarding its ability to pursue hydrogen bonds) which may imply a very different behaviour between the interaction of S and each of these surfaces. In order to complement our simulations of the interaction between S and the skin models, we will consider also simulations of the wetting behaviour of SC and sebum by placing a water droplet on top of each surface. As we will see, the results of these simulations are helpful in the interpretation of the results of the S - skin interaction simulations.

The organization of the paper is as follows. In Section II, we briefly describe the simulation methods. In Section III we discuss the composition and structure of the skin models considered in this work and we describe the results of simulations of the skin models in contact with a water droplet and with an hydrated spike S protein from SARS-CoV-2. In Section IV, we compare the obtained results with previous results in which we study the interaction of a solvated S protein with different materials. We finally end up with the Conclusions.

## 2 Simulation Methods

### 2.1 Description of Simulations

We have performed a total of four different simulations, as summarized in table 1 which correspond to wetting simulations of the two skin models (placement of a water droplet on top of each skin model) and their interaction with the S protein. In addition to these simulations, we have performed an additional simulation of the interaction of the spike protein with a POPC layer as a reference for the interaction between the protein and a lipid bilayer. The details and results for this case are given in the ESI.

**Table 1.** Summary of simulations performed in this work, indicating a label for each simulation, a description of the simulated system, the simulation box size, the total number of atoms and the total simulation time.

| Label | Simulated system | Box size (nm<sup>3</sup>) | Atoms | Sim time (ns) |
| --- | --- | --- | --- | --- |
| SC-W | SC and water droplet | 21.40 × 21.40 × 43.00 | 256706 | 30.9 |
| SC-S | SC and hydrated spike | 21.40 × 21.40 × 43.00 | 483473 | 49.2 |
| SB-W | Sebum and water droplet | 23.34 × 23.34 × 43.00 | 465835 | 29.7 |
| SB-S | Sebum and hydrated spike | 23.34 × 23.34 × 43.00 | 639733 | 50.2 |

All MD simulations were performed with NAMD 2.13 software^19^ using standard settings as described in the ESI.

The temperature was set at 305K in all simulations in order to reproduce physiological skin temperature. As in our previous work^14^, the force field employed in the simulations is the CHARMM36 force field, which includes parametrization of proteins, lipids and general organic molecules (CGenFF)^20^. This forcefield is, therefore, appropriate for describing both the spike glycoprotein and the systems considered in the paper. The water model used in our simulations was the TIP3P model included in CHARMM36.

The structures of the skin surfaces were prepared using CHARMM-GUI^21–24^, as described in the ESI. In the case of the stratum corneum (SC) surface we prepared a thermalized and equilibrated 1:1:1 mixture of cholesterol, ceramide CER-NS (24:0) and free fatty acid FFA (24:0) with an area of 21.4 × 21.4 nm^2^ and thickness of 9.46 nm. For the case of sebum, we prepared a mixture of triglyceride tri-cis-6-hexadecenoin, lauryl palmitoleate and squalene with a surface of 23.34 x 23.34 nm^2^ and a thickness of 8.9 nm. The chemical structures of these molecules are shown in Figure 2 and a snapshot of both equilibrated SC and sebum systems is shown in Figure 3.

**Fig. 2.**
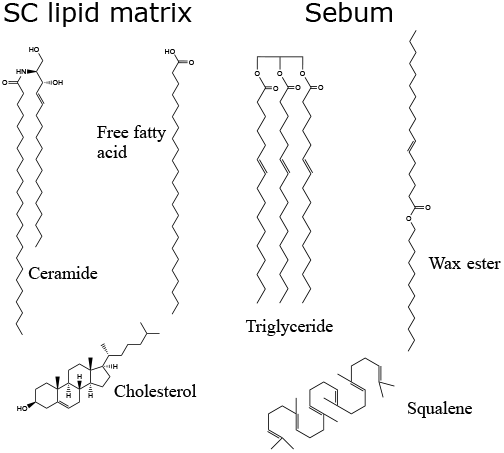
Molecules used in our model of the lipid matrix of the stratum corneum (SC) and sebum. The lipid matrix of SC is constituted by ceramides, free fatty acids and cholesterol. Sebum is constituted by triglycerides, wax esters and squalene.

**Fig. 3.**
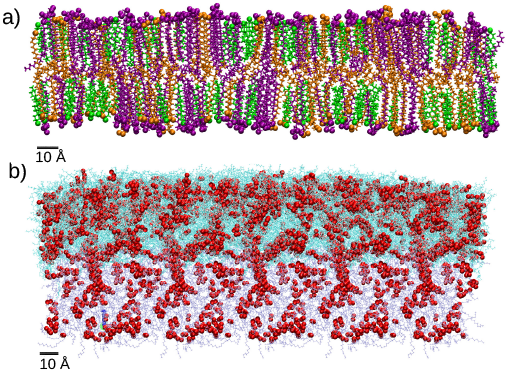
Snapshots of the two models for the skin surface used in our simulations. (a) Stratum corneum (SC) equilibrated bilayer with each molecule type shown in a different color: cholesterol (green), CER (purple) and FFA(orange). These molecules are shown in bond representations with but oxygen and nitrogen atoms are shown as spheres to indicate the location of the hydrophilic headgroups. (b) Equilibrated sebum layer. All molecules are shown as lines and oxygen atoms are highlighted as red spheres. The cyan color indicates free molecules whereas molecules in iceblue color are maintained in fixed positions during MD simulations. Oxygen atoms are represented as red spheres.

These two skin surfaces were later employed for the NVT simulations of wetting of skin by a water droplet (SC-W and SB-W in Table 1) and skin - S protein interaction (SC-S and SB-S in Table 1).

In the SC-W simulation, we consider the wetting of the SC lipids by a water droplet. To this end, we merge the equilibrated SC bilayer with a pre-equilibrated water droplet containing 622 water molecules. The water droplet was placed 4 Å next to the SC membrane. In SC-S simulation, we consider the interaction of the surface of SC lipids with the S protein spike. To this end, we merge the SC lipids structure with an equilibrated hydrated spike structure (obtained as described before) with the hydrated spike placed 10 Å from the SC surface.

In the SB-W simulation (wetting of sebum) we merge the obtained sebum structure with a pre-equilibrated water droplet constituted by 6845 molecules, situated 2 Å next to the sebum surface. In the SB-S simulation (SB - S protein interaction), we merge the sebum system structure with the one corresponding to the hydrated spike with the hydrated spike placed initially at 5 Å from the sebum surface.

In the case of SB-W and SB-S simulations, we recall that sebum has a fluid nature so it needs to be deposited onto an underlying surface. In simulations is not realistic to include any explicit surface, since it may perturb the sebum liquid structure. In real skin, the thickness of the sebum layer is large enough so that the outer sebum liquid is unperturbed. Therefore, in SB-W and SB-S simulations the atoms corresponding to the bottom part of the sebum were maintained at fixed positions during the MD simulation, as shown in Figure 3 (see also the ESI for details).

### 2.2 Analysis of Results

In the wetting simulations (SC-W and SB-W), we monitored the contacts between the water surface and each system, and the systems were considered equilibrated after 10 ns for the case of the SC lipids and 6 ns for the case of sebum when the droplet shape was completely stabilized. The simulations were later continued until a total time of ∼ 30 ns for both systems for production purposes. During production, we computed the 2-D water density profile using our own Fortran code, as in our previous work^25^. The contact angles were calculated by superimposing an equation of a straight line on plots of the water density and determining its slope using our own Python code.

In the case of SC-S and SB-S simulations we have calculated the evolution of RMSD, the number of contacts and the number of hydrogen bonds between the protein and the surface and the tilt angle of the protein with respect to the surface. The RMSD was computed between each instantaneous structure and the initial structure using the RMSD trajectory tool implemented in VMD^26^. In the calculation we considered all spike atoms except glycans and hydrogen atoms. The number of aminoacids in contact with each skin model was computed considering that a contact between aminoacids and skin occurs when at least one atom of the aminoacid is found at a distance smaller than 3.5 Å from any skin atom. In order to count the number of aminoacids at each time step, we employed a TCL script running on VMD implementing the distance requirement described above. The number of hydrogen bonds between the spike protein and skin molecules was computed using the build-in VMD plugin. We used an acceptor–donor distance cutoff of 3.5 Å and acceptor–hydrogen–donor angle cut-off of 30°. The tilt angle between the spike protein and the *z* axis (the axis perpendicular to the surface) was computed using a TCL script in VMD and it was defined as follows. The axis of the protein was taken following the the vector joining the center of mass of the protein and the center of mass of the residues 1070 to 1146 (which are two arbitrary residues located in an internal protein region far from the RBD of the protein). In order to calculate the proportion of each lipid in contact with the S protein we also used a 3.5 Å cutoff distance. For the calculation of average values, we considered as equilibrated configurations those from *t*=40 ns to the end of the simulation for both SC-S and SB-S.

All snapshots were made using Visual Molecular Dynamics (VMD) software.^26^

## 3 Results and Discussion

### 3.1 Modelling of the skin surface

As explained in the Introduction (see also Figure 1), we will consider two models for the skin surface, one corresponding to the top layer of a clean epidermial surface (the stratum corneum) and one corresponding to an oily skin (composed by sebum lipids).

The stratum corneum (SC) is the top layer of the epidermis, and its functions are mainly to limit the passive water loss, reduce the absorption of chemicals from the environment and prevent microbial infection ^17^. The structure of the SC is complex, but it can mainly be described with the so called “brick and mortar” model^17^. In this model, the “bricks” of the SC are cells called corneocytes (which have resistant cell envelopes made of cross-linked proteins instead of usual phospholipid membranes). The mortar are bilayer structures of lipids^17,27,28^ that cover the corneocytes “bricks” and are critical for the barrier function of the SC. Any external agent in contact with the SC will be in contact with the SC lipids, so in our simulations of the SC surface we will limit ourselves to considering a bilayer of SC lipids, as in previous simulations^27,29–31^. The SC lipids are a complex mixture of lipids of different types, so for simplicity we will consider here only the most abundant ones. We consider a 1:1:1 molar mixture of ceramides (CER), free fatty acids (FFA) and cholesterol (CHOL) as in Ref ^29^. Both CER and FFA are a wide class of lipids, with many different subclasses present in the stratum corneum^32,33^. The ceramide molecules consist of fatty acids amide linked to sphingoid bases, with long hydrocarbon chains with a broad distribution of carbon numbers and a small hydrophilic headgroup (smaller than those of phospholipids). In our model, we have selected the most common ceramide, which has the chemical structure shown in Figure 2. Concerning the free fatty acid, we selected also only the most abundant one (24 carbons), as shown in Figure 2. We consider the FFA molecule in its neutral (protonated state) as in previous simulations^29,30^. However, it is worth noting that the protonation state of FFA will depend on the local pH of the skin, which has a strong variability due to many factors (depending on factors such as water employed to clean skin, prior exposure to cosmetic products, atmosferic agents,…)^34^. Recent simulations take into account this variability in pH to asess the impact of pH on the interaction of external agents with skin ^31^.

As we mentioned before, the SC lipids are known to form a lamellar structure of assembled bilayers, so for the outer layer of SC lipids we consider a single bilayer (Figure 3a) build as detailed in the Methods section and the ESI.

As an “oily” skin surface model we consider a layer of sebum lipids, which covers several parts of human skin, as discussed in the Introduction. Sebum is a complex lipid mixture of triglycerides, was esters, squalene, free fatty acids and cholesterol esters ^18^ (see Table 2). In some previous simulation studies only one lipid component was included, either considering only triglycerides molecules^30^ or squalene^35^. Also, in Ref ^36^ the authors simulated a coarse-grain model in which eight different types of lipids were considered. Here, in our model of sebum we include each of the three main components: triglycerides, wax esters and squalene, with the molar fraction indicated in Table 2 which is similar to the actual biological composition. For triglycerides and wax esters (which are lipid types rather than single molecules), we select the most representative molecule (see Figure 2). As a representative triglyceride, we choose tri-cis-6-hexadecenoin which is the main triglyceride of human sebum together with the saturated equivalent lipid ^37^ and it has been used in previous sebum modeling studies ^30,38^. Concerning the wax esters, we chose lauryl palmitoleate because it is a representative sebum ester ^39^ but we should keep in mind that there is a huge variety both in length and degree of saturation of lipid chains in wax esters. Due to their chemical structure, the lipids of the sebum layer do not self-assemble into bilayers, they rather form an oily viscous liquid layer, as illustrated in Figure 3b.

**Table 2.** Composition of sebum according to experimental data from Ref^18^ and in our simulation model.

| Molecule | Molar proportion (%) |  |
| --- | --- | --- |
|  | Exp. data | Simulation model |
| Triglyceride | 45-50 | 56.8 |
| Wax ester | 25 | 29.7 |
| Squalene | 12 | 13.5 |
| FFA | 10 | 0 |
| Cholesterol ester | <5 | 0 |

### 3.2 Wetting behaviour of the Skin surface models

Our first study using the two skin surface models of Figure 3 is a characterization of their wetting behaviour by placing a water droplet on top, as described in the Methods section (simulations SC-W and SB-W in Table 1). The results are shown in Figure 4.

**Fig. 4.**
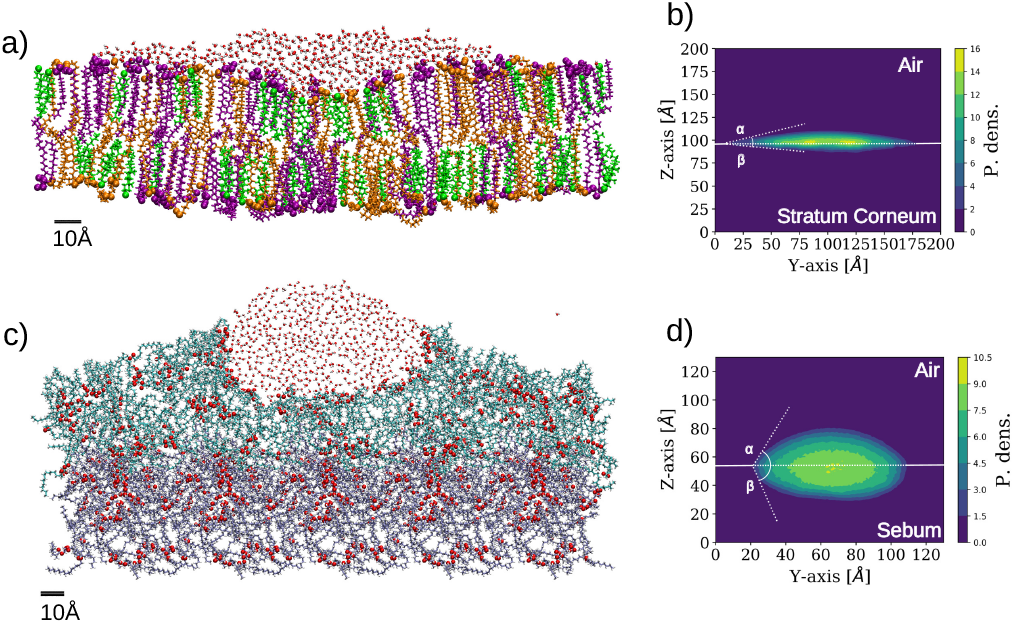
Simulation results for wetting of two different skin models. The images are made from a cut in the plane perpendicular to the surface to better distinguish the shape of the wetting region. The representation employed for the lipids is the same as employed in Figure 3. (a) Snapshot of SC-W simulation corresponding to the wetting of SC by a water droplet. (b) 2-D density profile of water molecules in SC-W simulation evaluated on a plane perpendicular to the SC bilayer (YZ). The two contact angles are indicated (*α* ≈17°, *β* ≈ 7°). (c) Snapshot of SB-W simulation corresponding to the wetting of sebum by a water droplet. (d) 2-D density profile of water molecules on plane YZ for the sebum surface. The two contact angles are indicated (*α* ≈ 63°, *β* ≈ 70°.)

Unlike wetting of a solid substrate, a droplet over a soft or liquid interface also deforms the interface at its contact and the droplet acquires a lens shape^40–42^. This is clearly observed in Figure 4 for both cases. In the case of wetting of soft surfaces, two contact angles can be defined, corresponding here to the three phase contact between skin, liquid water, and air (in fact, vacuum in our simulations) and the two phase contact between the skin and liquid water. These angles are designed as *α* and *β* in Figure 4, respectively. For the case of stratum corneum lipids (SC-W simulation), we obtained *α* =17° and *β* = 7° for the SC-water-air and SC-water interfaces, respectively (Figure 4b). In the case of sebum (SB-W simulation), these angles are *α* =63° and *β* =70° respectively as seen in Figure 4c. The differences between SC and sebum in terms of wetting properties are mainly due to the fact that the SC surface exposes a much larger amount of hydrophilic groups than sebum. This is not unexpected since the SC surface is a lipid bilayer exposing hydrophilic headgroups. In the case of sebum, these values for the contact angle correspond to an intermediate wettability of the sebum substrate by liquid water. In spite of the oily nature of sebum, there are chemical groups (the glycerol groups from the triglycerides) that are able to make hydrogen bonds with water, as highlighted in Figures 3b and 4c.

In this respect, it is interesting to compare the location of the oxygen atoms highlighted in the snapshots in Figure 3b (sebum) and Figure 4c (sebum with a water droplet on top). These figures suggest that sebum molecules rearrange around the water droplet in order to maximize exposure of lipid oxygen atoms to water, as expected from their hydrogen bonding capacity.

It is interesting to compare our simulations results with experimental measurements, keeping in mind that our simulations correspond to wetting by a nanoscopic water droplet of simplified models of skin whereas experimental data corresponds to macroscopic phenomena in heterogeneous and complex human samples.

Early experiments^43^ yielded an angle of 58° for water wetting of skin from a dorsal surface of a human finger. This value is similar to the value of *α* obtained in Figure 4d for sebum. The authors also repeated the measurements after washing hands with soap and water, rinsed with deonized water and dried. In this case, the angle increased to 104° demonstrating that the removal of lipid molecules by the use of personal healthcare products exposes an hydrophobic internal structure of skin (the underlying reason for a such high contact angle was not reported).

More modern measurements^44^ in skin from different parts of the body give similar values. For example, on forehead skin, a rich sebum site, this work reports contact angles about *θ*_*w*_ ∼ 57°–73°, which are again consistent with our results for sebum in Figure 4d. On the volar fore-arm, a poor sebum site, the measured contact angles^44^ were in the range *θ*_*w*_ ∼ 80°–90° so in this part of the body skin is more hydrophobic. Taken all together, these results are consistent with the view of skin lipids as a major factor in retaining skin hydration.

It is remarkable that, as we mentioned, our simulation results for wetting of our sebum model by a nanoscopic water droplet (SB-W simulation, Figure 4c,d) are consistent with the macroscopic contact angles measured for skin covered by sebum.

We also note that the results for the contact angle obtained in our simulations for the SC lipids model (Figure 4a,b) do not correspond to any experimentally measured contact angle. This can be interpreted as follows. As discussed before, we expect that in those parts of the skin in which sebum is not present, the stratum corneum should be directly exposed to the environment. The structure of the stratum corneum, following the previously mentioned “brick-and-mortar” model, is composed by corneocytes (not included in our simulations) embedded in a lipid matrix. The results discussed here suggest that the SC lipid matrix that embeds the corneocytes is not a determining factor of the macroscopic wetting properties measured experimentally.

### 3.3 Interaction with the Spike protein

The interaction between the Spike protein and the skin surface was studied in our simulations by initially placing the hydrated Spike protein near the sebum surface and the SC lipids surface (simulations SC-S and SB-S in Table 1). The details are described in the Methods section.

Snapshots corresponding to the initial and final (adsorbed) configurations for each case (Fig.5) indicate striking differences between the results for both simulations. In the case of adsorption onto the SC lipids (SC-S), the S protein adsorbs with its long axis almost parallel to the lipid surface whereas in the case of sebum (SB-S), the S protein adsorbs with an orientation close to its original perpendicular orientation (Fig.5). In this orientation the adsorbed Spike maintains the Receptor Binding Domain (RBD) of the three monomers of the spike oriented towards the surface.

**Fig. 5.**
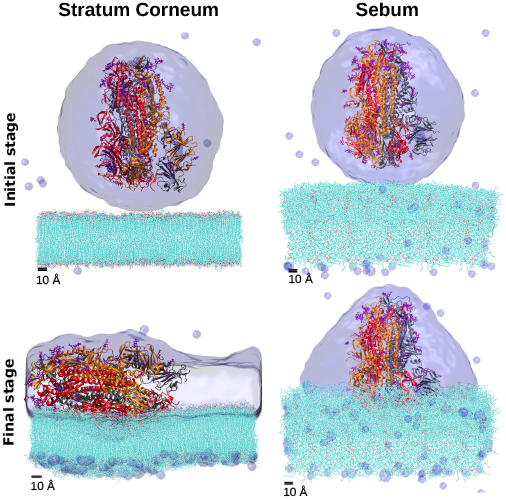
Snapshots of the MD simulations of the interaction between the Spike protein and SC and sebum surfaces (simulations SC-S and SB-S in Table1). We show the initial and final configurations (top and bottom, respectively). Hydration water is shown as a surface and the lipids in the skin models are shown as lines (with their oxygen atoms indicated in red). Each monomer of the trimeric spike is represented by a different color color (gray, orange, red) in cartoon representation.

As can be seen in these snapshots, the behaviour of the hydration water also differs in both cases. In the case of the S protein adsorbed onto sebum, S is maintained inside an hydration droplet, with a droplet shape compatible with the contact angle observed in the previously discussed wetting simulation SB-W. In the case of adsorption onto SC, we observe a full wetting of the SC bilayer. It is plausible that this wetting process may induce a tension over the protein, inducing its rotation over the SC surface.

A quantitative analysis of the evolution of protein adsorption in both cases (protein RMSD, the number of aminoacids in contact with lipids, the number of lipid-protein hydrogen bonds and the tilt angle) is given in Figure 6.

**Fig. 6.**
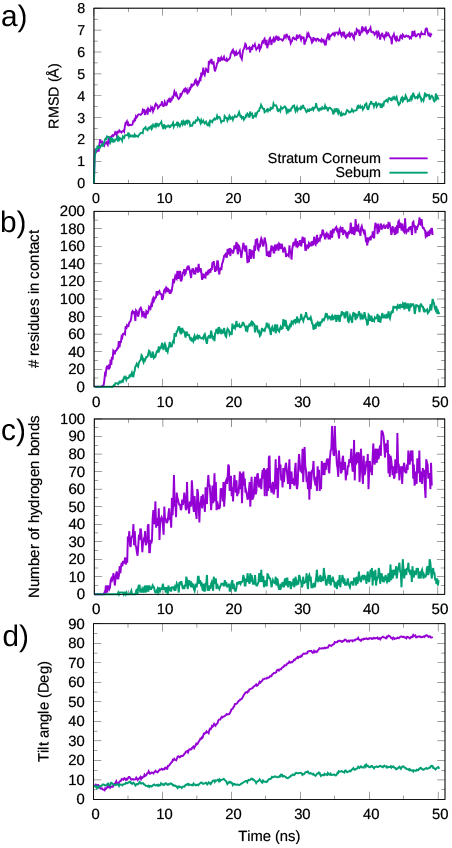
Time evolution of physical quantities in the MD simulations of the interaction between the Spike protein and SC and sebum surfaces (simulations SC-S and SB-S in Table 1). (a) RMSD as a function of time. (b) Number of S protein residues in contact with each skin surface as a function of time. (c) Number of hydrogen bonds between S protein and lipids as a function of time. (d) Tilt angle of the S protein respect the surface as a function of time. Purple lines correspond to the adsorption on the stratum corneum (simulation SC-S), while green lines correspond to adsorption on sebum (SB-S).

These results show that in the case of adsorption onto SC lipids (SC-S simulation), there is a continuous increase of all quantities over time from contact between the protein and the surface until reaching equilibrium values after ∼ 40 ns of simulation. According to Figure 6d, a final equilibrium angle of ∼83.0° was reached, corresponding to the near perpendicular orientation of the protein to the *z* axis (i.e. near parallel orientation respect to the surface) observed in the snapshot of Figure 5. In the case of adsorption onto sebum, the changes in all quantities were smaller and the system reaches equilibrium in a shorter time. For example, the protein orientation has only a modest increase over time, stabilizing at about 15° (see Fig. 6d), consistent with the snapshot of Figure 5.

In both cases there is a small change in the protein structure as measured by the RMSD (Figure 6a), which is larger in the case of SC lipids. In any case, both RMSD changes are smaller than those reported previously for adsorption of the Spike protein onto cellulose and graphite surfaces ^14^ (see discussion in section 4).

The number of S residues in contact with the skin surface (Figure 6b) is much larger for the case for S adsorption onto SC than for sebum, as can be expected from the larger surface of contact. But it is important to emphasize that even in the case of sebum the number of contacts is substantial. For comparison, it is larger (∼51 vs ∼87) than that observed for the interaction between S and a cellulose surface in our previous simulations^14^. A significant part of these contacts between S and skin lipids involve hydrogen bonds. According to Figure 6c, we have ∼70 protein-lipid hydrogen bonds which corresponds to ∼0.4 hydrogen bonds per aminoacid. In the case of sebum, the number of aminoacids in contact with surface molecules is about half the value obtained for SC lipids (∼ 90) and the number of hydrogen bonds is substantially smaller (∼ 10), as seen in Figure 6).

Given the lipid composition of the skin models, it is natural to ask whether all its component molecules will interact equally with the S protein (of course modulated by its proportion in the skin composition) or we can expect a substantially different interaction of different skin lipids with S. The analysis shown in Figure 7a indicates that for both the SC-S and SB-S simulations there is a preference of certain constituent lipids of the skin models with respect to others to interact with the spike protein.

**Fig. 7.**
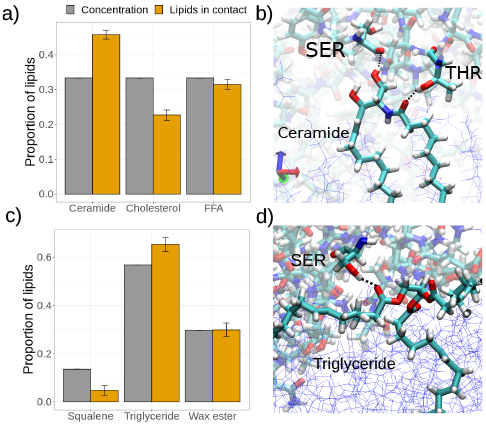
Analysis of the interaction between the S protein and skin lipids from the stratum corneum (panels (a) and (b)) and sebum (panels (c) and (d)). (a) Equilibrium proportion of lipids type in contact with the spike protein (orange) in comparison with the lipid concentration (gray). (b) Hydrogen bond detail between a ceramide and a serine and threonine residue of S protein. (c) Equilibrium proportion of lipids type in contact with the spike protein (orange) in comparison with the lipid concentration (gray). (d) Hydrogen bond detail between a triglyceride and a serine residue of S protein.

In the case of stratum corneum (simulation SC-S, Figure 7a) we have an equal molar fraction of the three components but the proportion of lipid in contact between each SC component and S protein does not follow this proportion. There is a clear excess of ceramide lipids in contact with S whereas there are less cholesterol molecules in contact with S than expected by the composition. A possible explanation is that a underlying mechanism for these contacts is hydrogen bonding. This is illustrated in Figure 7b, where we show a single ceramide lipid forming two hydrogen bonds with the S protein (with a serine and a threonine residue). Ceramide lipids have a bulky hydrophilic headgroup (see Figure 2) exposed at the interface with four available atoms for hydrogen bonds. Therefore, we can expect a strong affinity between ceramides and the S protein. On the contrary, cholesterol has a small hydrophilic headgroup made of a single -OH so it is expected to have a much weaker interaction with the S protein, as we obtain in Figure 7a.

In the case of sebum components (simulation SB-S, Figure 7c) we also observe a similar effect as observed previously with SC, namely there is a component with an excess proportion in contact with the protein (triglycerides) and a component with less proportion in contact with S than expected from the composition (squalene). This can also be interpreted with the help of the results of our previous SB-W simulation, in which we observed that oxygen atoms (mostly from triglycerides) in the sebum layer tend to go to the water interface after wetting (Figure 4b). Therefore, it is reasonable to expect now in the SB-S simulation that these oxygen atoms are able to make hydrogen bonds with the S protein, as illustrated in Figure 7d. On the contrary, the squalene molecule (see Figure 2) is unable to make hydrogen bonds, so it is expected to interact only weakly with the S protein.

It is also interesting to note that we have checked that the results obtained for the interaction between the spike protein and SC lipids do not dependent substantially on the particular lipid composition of the bilayer. To this end, we have performed an additional simulation considering the interaction of the spike protein and a pure POPC lipid bilayer (see ESI). POPC is a common phospholipid present in the animal cell membrane which assemblies in bilayers. The results are detailed in the ESI and also compiled in Table 3. They are quantitatively close to those obtained for the SC lipids, demonstrating that the exact composition of the bilayer is not influencing the results.

**Table 3.**
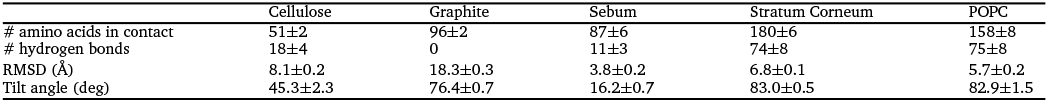
Comparison of results for MD simulations of the interaction of the SARS-CoV-2 S protein and different surfaces. The results for cellulose and graphite correspond to our previous work ^14^ whereas the results for stratum corneum, sebum and POPC correspond to the present work.

## 4 Interaction of S protein with different surfaces: A comparison of MD results

At this point it is interesting to compare our results for the spike - skin interaction (Figures 5 and 6) with our previous work^14^ corresponding to the interaction of the spike with cellulose and graphite. The results for all these systems are compiled in Table 3 and Figure 8. They correspond to results for four different surfaces, with a different hydrogen bonding capacity and in a different state or organization (polymer, solid, waxy liquid or self-assembled bilayer). Looking at these results, there are several interesting comparisons that can be made between these different materials.

**Fig. 8.**
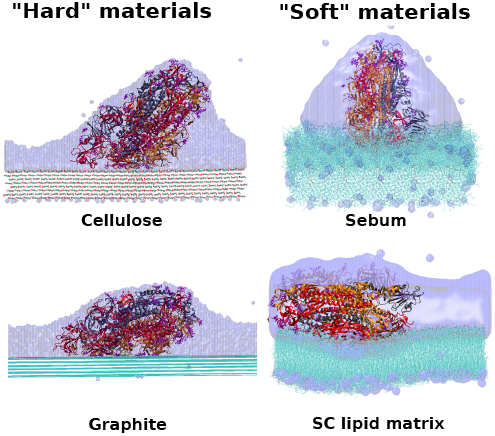
Comparison between hydrophobic-hydrophilic and hard-soft materials. Final stage for cellulose, graphite, SC and sebum surfaces.

Sebum is the surface that induces the smaller deformation in the S protein, retaining its original orientation while having a large number of sebum - protein contacts. In fact, sebum has a larger number of contacts with the protein than cellulose and a similar value (but slightly smaller) than graphite (see Table 3). Looking at the snapshots of Figure 8, we see that this is due to the fact that sebum is a soft material and the interface is able to deform to increase the contact with the spike. In the case of graphite, a similar number of contacts is obtained but with the cost of a substantial deformation of the protein. In fact, the case of graphite stands out as the one in which the deformation of the spike is larger (as measured by the RMSD, see Table 3).

The case with the larger number of hydrogen bonds much larger than any other case, (see Table 3) is that of the interaction of spike with stratum corneum lipids. The spike forms a slightly larger number of hydrogen bonds with cellulose than with sebum, although the numbers are similar. This similarity is remarkable since cellulose is a material with a high capacity to make hydrogen bonds^25^. As discussed in Section 3, this hydrogen bonding ability of sebum observed in simulations is due to the soft nature of sebum and the possibility of sebum molecules (mostly triglycerides) to re-organize and expose their oxygen atoms. In the case of cellulose there is also a larger deformation of the spike (as measured by the RMSD, see Table 3) than in the case of stratum corneum or sebum. It is also interesting to remark that as a general trend, we observe that in the case of hard surfaces (cellulose and graphite), the spike has a larger RMSD than in the case of soft surfaces (the two skin models). This seems to indicate that the ability of the soft interfaces to adapt to the incoming protein allows for a substantial interaction without the need of deformation of the spike protein.

## 5 Conclusions

In this work, we presented molecular dynamics simulations of the SARS-CoV-2 spike glycoprotein interacting with human skin. We have considered the interaction of spike with two different skin models, the lipid matrix of the stratum corneum (the most external layer of the epidermis) and sebum (present on top of the epidermis in certain parts of the body). In a simplified view of the skin, these models can be regarded as models of a “clean” skin and an “oily” skin, respectively.

Our simulation results show striking differences between the two cases. In the case of stratum corneum lipids, the spike protein adsorbs with its long axis almost parallel to the lipid surface, maximizing the contact between the spike and the stratum corneum surface. In the case of sebum, the spike protein adsorbs retaining its original perpendicular orientation, with the Receptor Binding Domain (RBD) of the three monomers of the spike oriented towards the sebum surface.

Interestingly, the behaviour of the hydration of the spike protein also differs in both cases. In the case of stratum corneum, we observe a full wetting of the SC bilayer which may compite with the tendency of S to remain hydrated, thus producing a tension that may affect the orientation of S over SC. In the case of the spike protein adsorbed onto sebum, S is maintained inside a well-defined hydration droplet formed on top of the sebum layer. These results are consistent with our simulation results for the wetting behaviour of both surfaces, in which we obtain a much smaller contact angle for a water droplet on top of stratum corneum as compared with sebum.

The spike protein has also a tendency to interact differently with the different molecules present in our stratum corneum and sebum models. We observe a stronger interaction of the spike (higher number of protein-molecule contacts) with those skin molecules with higher hydrogen bonding ability: ceramides in the case of stratum corneum and triglyrecides in the case of sebum.

The number of hydrogen bonds between the spike protein and skin molecules is much larger in the case of stratum corneum as compared with sebum. Also, it has to be noted that in the case of sebum, the number of hydrogen bonds with the spike protein is comparable with that obtained in previous simulations of the interaction between the spike and cellulose ^14^, which is a remarkable result given the high tendency of cellulose to form hydrogen bonds.

We do not observe a significant deformation or structural change of the spike protein after interaction with both skin models. This is in sharp contrast with our previous results obtained for the interaction between the spike protein and solid surfaces ^14^ (cellulose and graphite). The comparison of the results obtained for the different surfaces indicates that in the case of solid surfaces the protein tends to change in order to increase the interaction with the surface, whereas in the case of soft surfaces the optimization of the protein-surface interaction can be achieved by a small deformation of the surface. Our results indicate the decisive importance of the wetting and hydrogen bonding properties combined with their soft matter or solid character.

At this point, it is relevant to discuss the possible implications of our simulation results for the more complex question of the interaction of a SARS-CoV-2 virion with human skin and its possible implications for its transmission. Our results suggest that interactions with sebum will tend to maintain the integrity of the hydrated SARS-CoV-2 virus spike. Therefore, we expect that a virus particle deposited onto an oily skin will be able to retain the integrity and maintain its hydration due to the wetting properties of sebum. In the case of interaction with the stratum corneum lipids, it seems possible a disruption of the virus integrity as the S protein adsorbs to the surface maximizing the spike-SC lipids interactions.Of course, it is important to keep in mind the many limitations of our computational models and simulations. The most relevant is probably the fact that we are considering only the S1 subunit of the spike glycoprotein and not the entire virus envelope. This may affect the interaction of the S protein with a surface (particularly limiting their possible change in orientation and possible rupture between the S1 and S2 subunits). and it is also important in understanding how an may affect the integrity of the envelope. Another limitation is related to the complexity of skin surface. Some aspects are omitted from our model such as sweat, rugosity, porosity and protein content. In general, we can say that in our present study as well as in our previous work^14^ we considered the molecular detail of the interaction between the spike protein and surfaces, but important factors (full virial envelope structure and mechanical properties, surface nanostructure,…) operating at a larger scale are omitted. In order to incorporate these factors, work involving combination of our atomistic results and CG models of a full virion such as those recently developed by other groups^45^ is under way.

## Supporting information

Supplemental Information: technical details for simulations

## Acknowledgements

This work was supported by the Spanish Ministry of Science and Innovation through Grant No. RTI2018-096273-B-I00, the “Severo Ochoa” Grant No.CEX2019-000917-S for Centres of Excellence in R&D awarded to ICMAB and the FPI grant PRE2020-093689 awarded to M.D. We thank the CESGA supercomputing center for computer time and technical support at the Finisterrae supercomputer. M.D. is enrolled in the Material Sciences PhD program of the Universitat Autonoma de Barcelona.

---

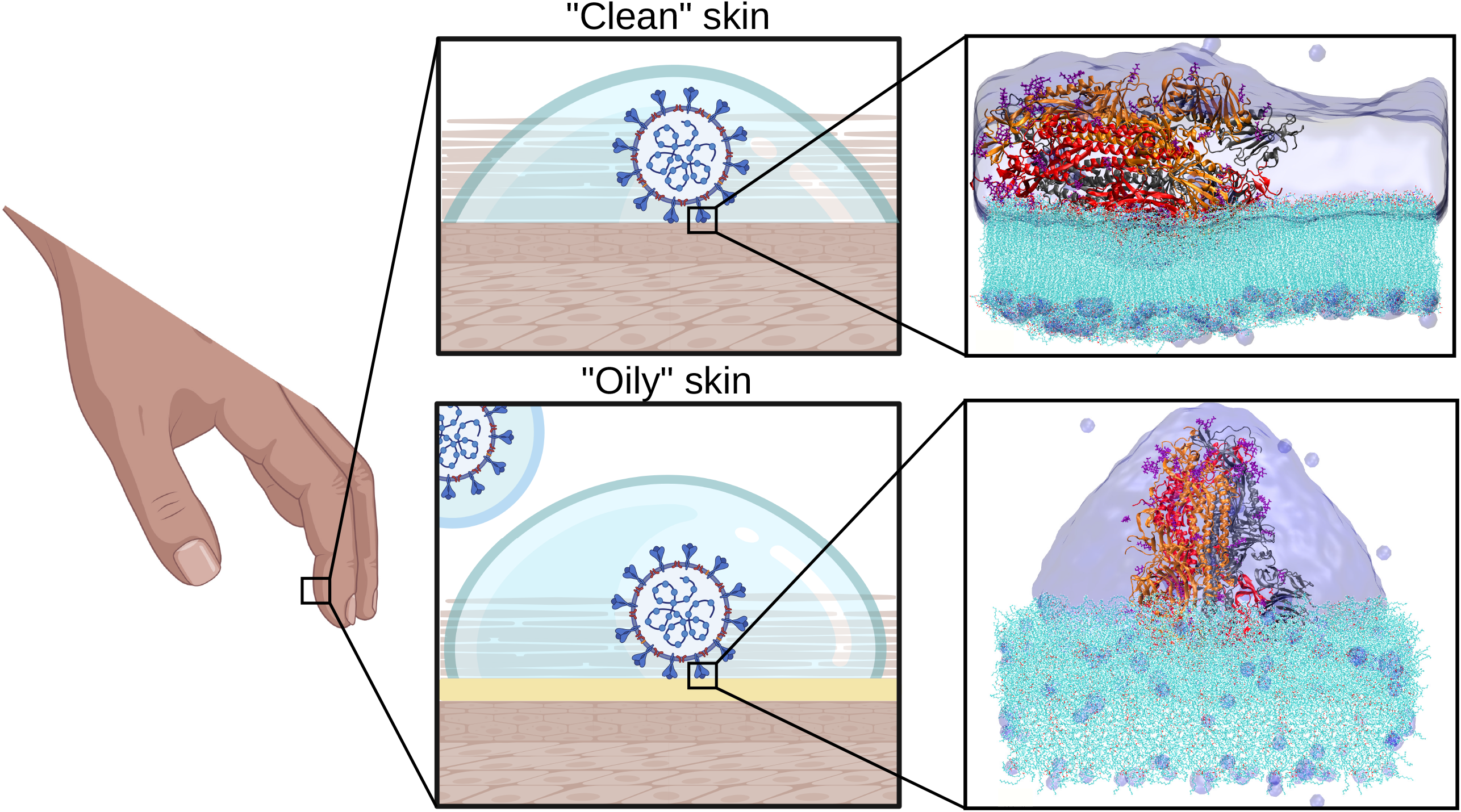

