## Supplemental Information: technical details for simulations for "Interaction between SARS-CoV-2 spike glycoprotein and human skin models: a molecular dynamics study"

### Electronic Supporting Information for "Interaction between SARS-CoV-2 spike glycoprotein and human skin models: a molecular dynamics study"

Marc Domingo and Jordi Faraudo*

#### 1 Further details for MD Simulations

##### 1.1 MD simulation parameters

In all simulations reported in Table 1 of the main paper the equations of motion were integrated with a 2 fs time step and electrostatic interactions were updated every 4 fs. All bonds between heavy atoms and hydrogen atoms were kept rigid. In all simulations we employed periodic boundary conditions in all directions. Lennard-Jones interactions were computed with a cutoff of 1.2 nm and switching function starting at 1.0 nm. Electrostatic interactions were computed using Particle Mesh Ewald (PME) algorithm using a real space cutoff set at 1.2 nm and a PME grid at 1.0 Å.

The temperature was set at 305K in all simulations employing the Langevin thermostat with a damping coefficient of 1 ps $^{-1}$. In NPT simulations we employed a Nosé-Hoover isotropic barostat with an oscillation period of of 50 fs and a damping time of 25 fs.

##### 1.2 Preparation and Equilibration of the SC bilayer model

An atomistic model for a single bilayer membrane of the stratum corneum (SC) lipids with equimolar composition was built using the Input Generator module Membrane Builder [1] of CHARMM-GUI [2–4]. The initial SC structure had a surface with dimensions 24.01 x 24.01 $nm^2$ and a thickness of 8.5 nm including hydration water molecules. This structure was equilibrated following the standard CHARMM-GUI protocols of minimization and thermalization. A further equilibration NPT run was performed, until the bilayer area was stabilized, requiring a total of 10.73 ns. In order to obtain a SC bilayer structure suitable for our subsequent simulations, we remove all the hydration water molecules from the final configuration. The obtained dry bilayer is shown in Figure 3a and it was used as starting point for two subsequent simulations involving wetting of SC and interaction of SC with S protein, SC-W and SC-S as described in the Methods section of the main paper.

##### 1.3 Preparation and Equilibration of the sebum layer model

The force field parameters for the three kinds of lipids molecules conforming the sebum (see main paper) were obtained using the Input Generator module Ligand Reader and Modeler [5] of CHARMM-GUI. Coordinate, topology and parameters files employed for these three sebum lipids are available for download [6]. An initial bulk system for the simulation of sebum was build using the "Input Generator" module of CHARMM-GUI. The initial sebum structure is a cubic box of 5 nm in length containing 42 molecules of triglyceride tri-cis-6-hexadecenoin, 22 molecules of lauryl palmitoleate and 10 molecules of squalene. The system was minimized and thermalized following the standard CHARMM-GUI protocols. The resulting bulk sebum structure had a liquid-like appearance in sharp contrast with the bilayer structure adopted by the SC lipids. In order to obtain a sebum system with the correct density, a further NPT simulation was performed with a total of 110 ns. The density of the obtained sebum system is 882±5 kg·$m^{-3}$. This value is slightly smaller than that of other previous MD sebum models [7] because in our sebum composition we included squalene (absent in the previous model) which has a smaller density.

The resulting equilibrated sebum system was too small for our subsequent simulations, particularly for the adsorption of the S protein, so we constructed a bigger system by replicating its molecules in the three directions. The dimensions of the resulting sebum surface (shown in Figure 3b) were $23.34 \times 23.34 \times 8.9 \text{nm}^3$.

##### 1.4 Preparation and equilibration of the hydrated Spike protein

The atomic coordinates for the SARS-CoV-2 spike glycoprotein structure were obtained from a cryo-EM structure [8] solved at 3.46 Å average resolution (PDB ID: 6VSB) as in our previous work [9]. This structure contains S1 and S2 spike subunits (with one RBD in "up" conformation) and a glycosylation pattern characterized by N-acetyl-D-glucosamine(NAG) residues. The only modification made to this initial structure was the addition with VMD of missing hydrogen atoms and connecting links between the protein amino acids and the NAG residues. The obtained structure contains 46 708 atoms and its total charge (assuming pH 7) is -23e.

The spike structure was solvated using VMD with an spherical solvation shell in order to maintain its hydrated functional state. The number of TIP3P water molecules added to solvate the glycoprotein was 60 634. We also added 23 $Na^+$ counterions to neutralize the charge of the spike. The system made by the hydrated spike with counterions has a total of 228 633 atoms. Finally, the hydrated spike system was thermalized as in our previous work [9] and the final structure was ready to be employed in the SC-S and SB-S simulations.

##### 1.5 Preparation and Equilibration of the POPC bilayer model

Alternatively, atomistic simulations of a POPC bilayer were made in order to show whether the interaction of the S protein with the SC lipid matrix is analogous to other bilayers.

The POPC bilayer was prepared using the Input Generator module Membrane Builder of CHARMM-GUI. The initial POPC

*Institut de Ciencia de Materials de Barcelona (ICMAB-CSIC), Campus de la UAB, E-08193 Bellaterra, Barcelona, Spain.*

structure had the same dimensions than the initial SC structure (a surface with dimensions 24.01 x 24.01 $nm^2$ and a thickness of 8.5 nm including hydration water molecules). The constructed system contains 844 POPC molecules per leaflet.

As in the case of SC lipid system, this structure was equilibrated following the standard CHARMM-GUI protocols of minimization and thermalization. A further equilibration NPT run was performed, until the bilayer area was stabilized, requiring a total of 10.0 ns. In order to obtain a SC bilayer structure suitable for our subsequent simulations, we remove all the hydration water molecules from the final configuration.

#### 2 Results for POPC system

Figure 1 shows the initial and final configuration of the spike glycoprotein onto a POPC bilayer. The similarities with the snapshots reported for the SC case in the main paper are clear. In Figures 2, 3 and 4 we report the same quantitative analysis for the spike - POPC bilayer as done in the main paper (RMSD, number of contacts and number of hydrogen bonds). For the sake of comparison, we also include the data for sebum and SC bilayer also reported in the main paper. The similarities between the POPC and SC results are clear, indicating the spike - lipid bilayer interaction does not significantly depends on the exact composition of the bilayer.

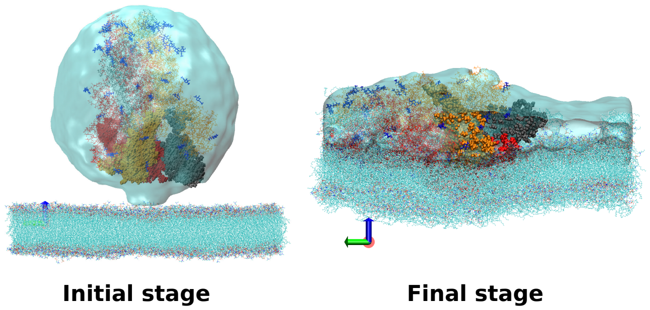


**Fig. 1** Initial and final configuration of POPC bilayer and spike protein.

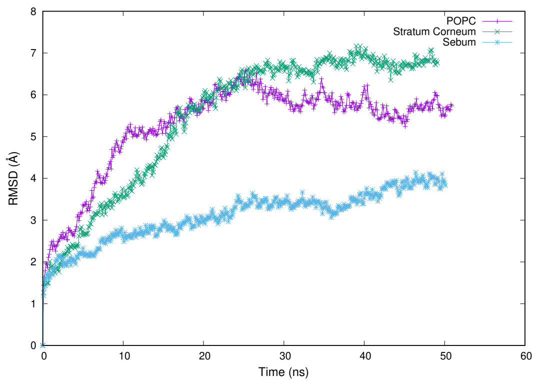


**Fig. 2** RMSD for POPC simulation in comparison with SC lipids and sebum system.

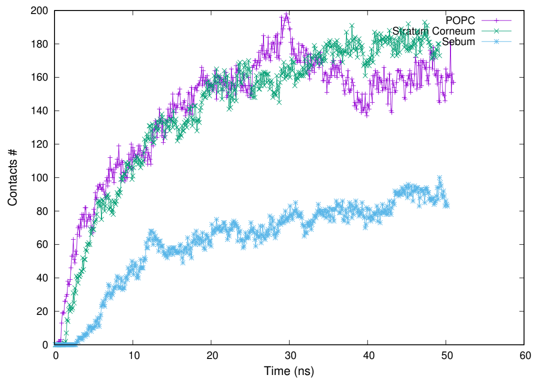


**Fig. 3** Evolution of the number of contacts for POPC simulation in comparison with SC lipids and sebum system.

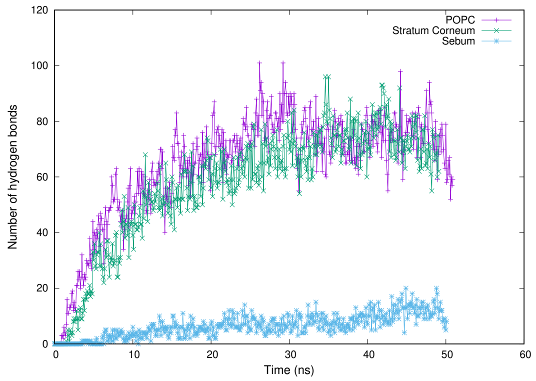


**Fig. 4** Evolution of the number of hydrogen bonds for POPC simulation in comparison with SC lipids and sebum system.

#### Notes and references


1 E. L. Wu, X. Cheng, S. Jo, H. Rui, K. C. Song, E. M. Dávila-Contreras, Y. Qi, J. Lee, V. Monje-Galvan, R. M. Venable, J. B. Klauda and W. Im, *Journal of Computational Chemistry*, 2014, **35**, 1997–2004.

2 S. Jo, T. Kim, V. G. Iyer and W. Im, *Journal of Computational Chemistry*, 2008, **29**, 1859–1865.

3 B. R. Brooks, C. L. Brooks, A. D. Mackerell, L. Nilsson, R. J. Petrella, B. Roux, Y. Won, G. Archontis, C. Bartels, S. Boresch, A. Caflisch, L. Caves, Q. Cui, A. R. Dinner, M. Feig, S. Fischer, J. Gao, M. Hodoscek, W. Im, K. Kuczera, T. Lazaridis, J. Ma, V. Ovchinnikov, E. Paci, R. W. Pastor, C. B. Post, J. Z. Pu, M. Schaefer, B. Tidor, R. M. Venable, H. L. Woodcock, X. Wu, W. Yang, D. M. York and M. Karplus, *Journal of Computational Chemistry*, 2009, **30**, 1545–1614.

4 J. Lee, X. Cheng, J. M. Swails, M. S. Yeom, P. K. Eastman, J. A. Lemkul, S. Wei, J. Buckner, J. C. Jeong, Y. Qi, S. Jo, V. S. Pande, D. A. Case, C. L. Brooks, A. D. MacKerell, J. B. Klauda and W. Im, *Journal of Chemical Theory and Computation*, 2016, **12**, 405–413.

5 S. Kim, J. Lee, S. Jo, C. L. Brooks, H. S. Lee and W. Im, *Journal of Computational Chemistry*, 2017, **38**, 1879–1886.

6 M. Domingo and J. Faraudo, *Structure and coordinate files for MD simulation of sebum lipids*, 2020, `https://github.com/soft-matter-theory-at-icmab-csic`.

7 A. S. Tascini, M. G. Noro, R. Chen, J. M. Seddon and F. Bresme, *Physical Chemistry Chemical Physics*, 2018, **20**, 1848–1860.

8 D. Wrapp, N. Wang, K. S. Corbett, J. A. Goldsmith, C.-L. Hsieh, O. Abiona, B. S. Graham and J. S. McLellan, *Science*, 2020, **367**, 1260–1263.

9 D. C. Malaspina and J. Faraudo, *Biointerphases*, 2020, **15**, 051008.
